# Removal of KCNQ2 from Parvalbumin-expressing Interneurons Improves Anti-Seizure Efficacy of Retigabine

**DOI:** 10.1101/2020.12.09.417295

**Authors:** Junzhan Jing, Corrinne Dunbar, Alina Sonesra, Ana Chavez, Suhyeorn Park, Heun Soh, Maxwell Lee, Anastasios Tzingounis, Edward C. Cooper, Xiaolong Jiang, Atul Maheshwari

## Abstract

Anti-seizure drug (ASD) targets are widely expressed in both excitatory and inhibitory neurons. It remains unknown if the action of an ASD upon inhibitory neurons could counteract its beneficial effects on excitatory neurons (or vice versa), thereby reducing the efficacy of the ASD. Here, we examine whether the efficacy of the ASD retigabine (RTG) is altered after removal of the Kv7 potassium channel subunit KCNQ2, one of its drug targets, from parvalbumin-expressing interneurons (PV-INs). Parvalbumin-Cre (PV-Cre) mice were crossed with *Kcnq2*-floxed (*Kcnq2*^fl/fl^) mice to conditionally delete *Kcnq2* from PV-INs. In these conditional knockout mice (cKO, PV-*Kcnq2*^fl/fl^), RTG (10 mg/kg, i.p.) significantly delayed the onset of either picrotoxin (PTX, 10 mg/kg, i.p)- or kainic acid (KA, 30mg/kg, i.p.)-induced convulsive seizures compared to vehicle, while RTG was not effective in wild-type littermates (WT). Immunostaining for KCNQ2 and KCNQ3 revealed that both subunits were enriched at axon initial segments (AISs) of hippocampal CA1 PV-INs, and their specific expression was selectively abolished in cKO mice. Accordingly, the M-currents recorded from CA1 PV-INs and their sensitivity to RTG were significantly reduced in cKO mice. While the ability of RTG to suppress CA1 excitatory neurons in hippocampal slices was unchanged in cKO mice, its suppressive effect on the spike activity of CA1 PV-INs was significantly reduced compared with WT mice. In addition, the RTG-induced suppression on intrinsic membrane excitability of PV-INs in WT mice was significantly reduced in cKO mice. These findings suggest that preventing RTG from suppressing PV-INs improves its anticonvulsant effect.

**Significance Statement:** One out of three patients with epilepsy remain resistant to anti-seizure drugs (ASD). Therefore, strategies that can enhance the efficacy of current ASDs or aid in the development of novel ASDs are in high demand. Here we show that by sparing a subset of inhibitory neurons, the ASD retigabine becomes more efficacious. These findings indicate that developing the ASDs that have relatively greater effects on excitatory over inhibitory neurons may be an effective strategy to improve the efficacy of ASDs.

## Introduction

The primary targets of most anti-seizure drugs (ASDs) generally fall into four major categories: (1) voltage-gated sodium channels; (2) voltage-gated calcium channels; (3) ionotropic receptors for neurotransmitters (γ-aminobutyric acid and glutamate); and (4) newer molecular targets including the synaptic vesicle protein SV2A and Kv7/KCNQ potassium channels (Rogawski, 2013). It is well known that these drug targets are not specifically confined to excitatory neurons; instead, they are widely expressed across distinct cell types in numerous brain regions, including inhibitory interneurons. Despite constituting a relatively small neuronal population, inhibitory interneurons exert diverse, powerful modulatory influences on excitatory neurons and circuit behavior. It remains an open question whether ASD action upon inhibitory neurons may counteract their beneficial effects on excitatory neurons (or vice versa), thereby reducing the efficacy of the ASD.

Retigabine (RTG) is a first-in-class ASD, and the primary mechanism for its anti-seizure effect is to suppress neuronal excitability by promoting the open state of a subfamily of voltage-gated potassium channels formed by KCNQ2-KCNQ5 subunits (Kv7 family), encoded by *Kcnq2*-*Kcnq5* (Main et al., 2000; Gunthorpe et al., 2012). Kv7 potassium channels are expressed in both excitatory neurons and GABAergic interneurons, including parvalbumin-expressing interneurons (PV-INs) (Cooper et al., 2001; Lawrence et al., 2006; Fidzinski et al., 2015). To directly examine how RTG’s simultaneous effects on inhibitory interneurons contribute to its overall efficacy, we selectively deleted *Kcnq2* from PV-INs (Soh et al., 2014), a major population of GABAergic inhibitory interneurons in the cortex (DeFelipe et al., 2013). We then tested whether the sensitivity to RTG in suppressing chemoconvulsant-induced seizures could be altered in these conditional KO mice (cKO). We also determined how the effects of RTG on intrinsic and evoked firing properties of both excitatory neurons and PV-INs in the hippocampus CA1 area were changed in cKO mice *in vitro*. While RTG was not effective in preventing seizures in two models of chemoconvulsant-induced seizures in wild type mice (WT), we found that RTG became effective in preventing seizures in cKO mice. In vitro, RTG suppressed the excitability of both pyramidal neurons and PV-INs in WT mice, but their effects on PVs-INs were selectively decreased in cKO mice where *Kcnq2* was conditionally removed from PV-INs. These findings suggest that RTG was more effective in preventing seizures when its suppressive effects on inhibitory neurons were selectively blunted, warranting the development of cell-type specific ASDs as an important avenue for future investigation.

## Materials and Methods

### Animals

All experiments were performed according to the guidelines of the Institutional Animal Care and Use Committee of Baylor College of Medicine. Parvalbumin-Cre (PV-Cre)/Ai9 mice (crossing PV-Cre mice with Ai9 stop-floxed mice, Jackson Laboratory Stock #008069 and #007909, respectively), a line shown to have recombination in >90% of cells expressing parvalbumin (Hippenmeyer et al., 2005), were crossed with *Kcnq2*-floxed mice (*Kcnq2*^fl/fl^) (Soh et al., 2014) to generate conditional knockout (cKO, PV-*Kcnq2*^fl/fl^) mice with PV-INs labeled with the red fluorescent protein tdTomato on a C57BL6/J background. Mice were group-housed in cages with a natural light-dark cycle and access to a standard pellet diet and water ad libitum. All mice were between postnatal day (P) 30-45 at the time of experiments. Genotype was blinded to investigators prior to analysis. For all experiments, both male and female mice were used, with no significant differences found between groups; therefore, data was pooled together. If necessary, mice were euthanized by CO_2_ inhalation (SmartBox, Euthanex, Palmer, PA).

### In vivo EEG monitoring

Mice were treated with meloxicam (5 mg/kg, Henry Schein Veterinary) and local injection of 50 µL of 2% lidocaine (Henry Schein Veterinary) before being anesthetized with isoflurane (2-4% in O_2_; Henry Schein Veterinary). Mice were surgically implanted with bilateral epidural silver wire electrodes over the somatosensory cortex (1 mm posterior and 3 mm lateral to bregma) and frontal cortex (1 mm anterior and 1 mm lateral to bregma) (Maheshwari et al., 2013). Reference and ground electrodes were placed over the left and right cerebellum, respectively. Implanted mice received daily meloxicam (5 mg/kg) injections for 3 days following surgery. Mice were allowed to recover for at least 2 days prior to recording (Soh et al., 2018). Freely moving mice were video-EEG monitored (AD Instruments), and power was analyzed and compared between groups as previously described, with left and right frontal electrodes re-referenced to left and right parietal electrodes, respectively (Maheshwari et al., 2017). Power between hemispheres was then averaged before comparing between WT and cKO mice.

### Drugs

Kainic Acid (KA; Santa Cruz Biotechnology) was dissolved in phosphate-buffered saline (PBS; Gibco, Life Technologies), and retigabine (RTG; Alomone Labs, Inc) and picrotoxin (PTX; Tocris) were first dissolved in dimethyl sulfoxide (DMSO) prior to being diluted to 1% volume/weight in PBS.

### Seizure Monitoring

Mice were treated with RTG 15 minutes prior to PTX or KA injection (RTG and PTX at 10 mg/kg; KA at 30mg/kg, i.p.), starting between 9:00-11:30 am to prevent confounding diurnal variation. Seizures were defined as rearing and falling with forelimb clonus (Racine Stage 5) (Makinson et al., 2014; Soh et al., 2014). For the PTX experiments, control mice either received no injection or received the vehicle injection (DMSO in PBS) prior to PTX administration. There was no significant difference in induced seizure latency between two cohorts of mice (vehicle injection (*n*=21) vs no injection (*n*=38)), so these mice were grouped together as ‘control’ mice. For the KA experiments, control mice received the injection of PBS alone.

### Behavioral Testing

Three behavioral tests were performed over a 3-day period. Mice were randomized to perform either the Tail Suspension Test or the Heat Sensitivity Test on Day 1, followed by the alternate test on Day 2. Mice underwent training for the Balance Beam Assay on both Day 1 and Day 2 prior to final testing on Day 3 as detailed below. The Tail Suspension Test was utilized to quantify depressive behaviors as previously described (Can et al., 2012). Briefly, the tail was attached to a suspension bar at a height of 50 cm for six minutes. The total time spent mobile or exhibiting escape behaviors was recorded. The Heat Sensitivity Test was utilized to assess response to painful stimuli (Moqrich et al., 2005; Deuis et al., 2017). Unrestrained mice were placed on a heating pad at 55°C and the time until onset of nocifensive behaviors was recorded. Finally, the Balance Beam Assay was utilized to evaluate motor balance and coordination (Luong et al., 2011). A flat, plexiglass beam (12.7 mm x 6.35 mm x 90 cm) was placed at 50 cm elevation with an aversive light stimulus placed atop the beginning end of the beam. During the first two days of training, each mouse crossed both the 12.7 mm width beam and 6.35 mm width beam three times. On the third day, the time taken to cross the center 75 cm of the beam during the two continuous crossings was determined and averaged (Luong et al., 2011). Comparison between groups for behavioral tests was performed with the Mann-Whitney U-test.

### Immunohistochemistry

For KCNQ2 detection in fluorescent-positive PV-INs (expressing tdTomato), we followed a modified fluorescence immunohistochemistry (IHC) protocol (Battefeld et al., 2014). Of note, 2% paraformaldehyde (PFA) was empirically found to be optimal in preserving both tdTomato and Kcnq2 immunostaining signals; higher concentrations of PFA limited the detection of KCNQ2 staining, while lower concentrations limited the detection of tdTomato staining. Mice were perfused transcardially with 20 mL PBS followed by 20 mL of 2% PFA. The brain was then removed, post-fixed in 2% PFA for 30 minutes and then suspended in 30% sucrose in PBS overnight. 20 µm sagittal sections were then cut on a cryotome, incubated first overnight with mouse anti-ankyrin-G (AnkG) (Rasband et al., 1999) (Sigma, #S8809) and rabbit anti-KCNQ2 (Cooper et al., 2001) as primary antibodies, followed by Alexa-Fluor goat anti-mouse 647 and goat anti-rabbit Alexa-Fluor 594 (Life Technologies) as secondary antibodies for one hour. Slices were then incubated in Prolong Gold Antifade with 4′,6-diamidino-2-phenylindole (DAPI) prior to imaging with either wide-field or confocal microscopy. Co-localization between KCNQ2 and AnkG signals was quantified by identifying AIS segments with AnkG expression for at least 20 µm (Höfflin et al., 2017) with either the presence or absence of coexpression with KCNQ2.

To detect KCNQ3 in PV-INs, a different IHC protocol was used given poor visualization of KCNQ3 with its antibody on 2% PFA-fixed tissue, as previously described (Pan et al., 2006; Battefeld et al., 2014). Briefly, PV-INs were filled with Alexa 488 during whole-cell recording. After the recording, the brain slices were immediately fixed with methanol at −20 °C for 20 minutes. Slices were then washed with PBS, blocked with non-fat milk, and incubated overnight with mouse anti-AnkG (Sigma, #S8809) and guinea pig anti-KCNQ3 (Devaux et al., 2004) as primary antibodies, followed by Alexa-Fluor goat anti-mouse 647 and goat anti-guinea pig Alexa-Fluor 594 (Life Technologies) as secondary antibodies for one hour. Slices were then incubated in DAPI prior to imaging with either wide-field or confocal microscopy. For all IHC studies, at least 4 mice from each genotype were used.

### Slice Preparation and Electrophysiology

Slice preparation from the mouse hippocampus followed an N-Methyl-D-glucamine (NMDG) slicing protocol (Ting et al., 2014; Jiang et al., 2015). Briefly, mice between P30-33 were deeply anesthetized using 3% isoflurane and immediately decapitated. The brain was then immediately removed from the skull and placed into a cold (0−4 °C) oxygenated NMDG solution. Parasagittal slices (300 µm) were cut with a vibratome (VT1200, Leica). The slices were incubated at 34.0±0.5 °C in oxygenated NMDG solution for 10-15 minutes before being transferred to an artificial cerebrospinal fluid solution (ACSF) containing: 125 mM NaCl, 2.5 mM KCl, 1.25 mM NaH_2_PO_4_, 25 mM NaHCO_3_, 1 mM MgCl_2_, 25 mM glucose and 2 mM CaCl_2_ (pH 7.4) for about 1 h. The slices were then incubated at room temperature for ∼0.5−1 h before recording.

Whole-cell recordings were performed as described previously (Jiang et al., 2013, 2015; Robbins et al., 2013; Greene et al., 2017; Varghese et al., 2021). Briefly, patch pipettes (5−7 MΩ) were filled with an internal solution containing 120 mM potassium gluconate, 10 mM HEPES, 4 mM KCl, 3 mM EGTA, 4 mM MgATP, 0.3 mM Na_3_GTP, 10 mM sodium phosphocreatine and 0.5% biocytin (pH 7.25). Whole-cell recordings of CA1 neurons were performed using Quadro EPC 10 amplifiers (HEKA). In many cases, multiple cells were recorded simultaneously (multi-patch recording) in the same slice to improve efficiency in data collection (Jiang et al., 2013, 2015). The spiking activity of each neuron in response to increasing current steps or ramp current injections (120 pA/s) was recorded with or without bath application of RTG (10 µM). Recorded neurons were filled with biocytin and then fixed for post hoc morphological recovery to further confirm cell identity (Jiang et al., 2015).

We followed a standard protocol to isolate M-currents (*I*_M_) from hippocampal neurons (Shah et al., 2002; Lawrence et al., 2006; Soh et al., 2014). Briefly, cells were first clamped at −20 mV, and then a 1 s long, −30 mV step was introduced every 15 s (Fig. 5). The *I*_M_ amplitude was measured from the deactivation relaxation at −50 mV (Fig. 5) (Soh et al., 2014). After obtaining a 5 min baseline, the Kv7 channels agonist retigabine (RTG) (10 µM) was added to the bath and the changes in *I*_M_ and the holding current (*I*_hold_) were recorded.

## Results

### *Increased anti-seizure efficacy of RTG in conditional KO mice (*cKO, PV*-Kcnq2*^fl/fl^)

Conditional KO mice (cKO, PV*-Kcnq2*^fl/fl^) exhibited normal outward appearance and growth throughout development (Fig. 1A, B). Spontaneous seizures were never observed in either genotype. Mortality up to 6 weeks of age was also not different between genotypes (Fig. 1C). WT and cKO mice had no significant difference in EEG spectral power (n=5 for each group, Fig. 1D) recorded over bilateral frontoparietal regions. Finally, there were no differences in performance on the tail suspension, heat sensitivity, and balance beam tests (n=9-12 per group, *p*>0.05, Fig. 1E-G). Altogether, these findings demonstrate that the baseline phenotype of cKO mice was largely unchanged.

**Figure 1.**
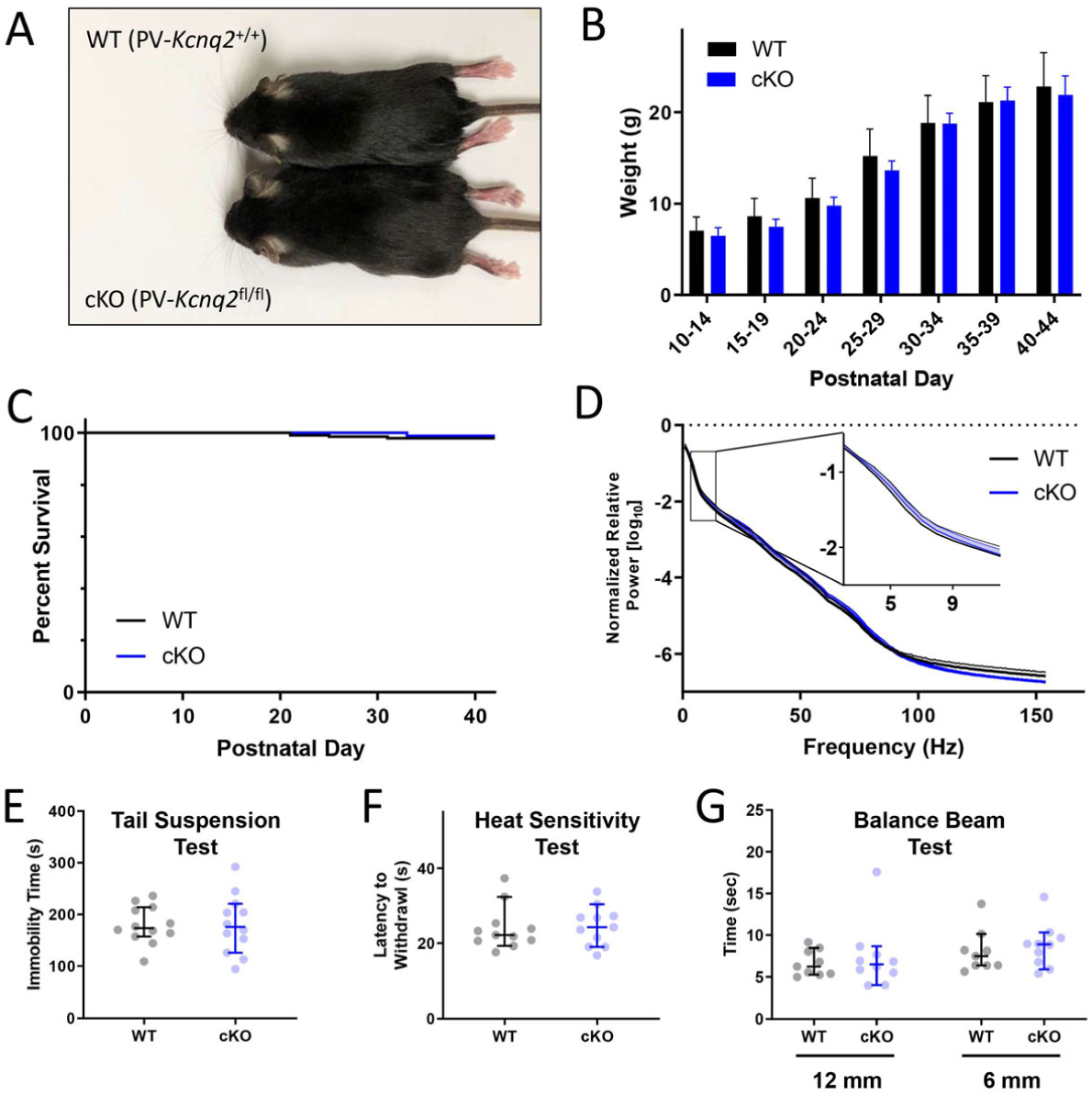
Characterization of cKO (PV-*Kcnq2*fl/fl) mice. (A) Representative WT (PV-*Kcnq2*+/+) and littermate cKO mice at P30. (B) No difference in relative growth curves across genotypes (n= 18,10). (C) Survival rate of WT (n=163) and cKO mice (n=101, *p*>0.05, Mantel-Cox test) as a function of postnatal day. (D) No difference between genotypes in EEG spectral power at frequencies between 2-150 Hz (n=5, mean±SEM, *p*>0.05, Tukey’s multiple comparisons test). There was also no significant difference between groups in performance on the (E) Tail Suspension Test (n=12 each, median±95% confidence intervals, *p*=0.887, Mann-Whitney U-test), the (F) Heat Sensitivity Test (n=11 each, *p*=0.699), or the (G) Balance Beam Test (n=9-10, *p*=0.434)

We first used a picrotoxin (PTX) seizure induction model to test if the anti-seizure efficacy of retigabine (RTG), an ASD targeting KCNQ2-containing channels, could be altered in cKO mice where KCNQ2 was selectively removed from PV-expressing interneurons (PV-INs), a major group of GABAergic interneurons. PTX-induced seizure semiology in cKO mice was similar to that seen in their WT littermates, including behavioral progression to characteristic Racine Stage 5 generalized convulsions (Soh et al., 2018). cKO mice had a shorter latency to seizure onset with PTX challenge than WT littermates (Fig. 2A,2B), as seen previously (Soh et al., 2018). Administration of RTG (10 mg/kg, i.p.) 15 minutes before PTX injection resulted in no significant difference in latency to seizure onset in WT mice when compared to control (*p*=0.405, Gehan-Breslow-Wilcoxon test, Fig. 2A). In contrast, RTG pretreatment in cKO mice significantly delayed the median time to seizure onset after PTX from 8.12 minutes to 14.51 minutes (190.2%; *p*=0.003, Fig. 2B), indicating that the selective removal of KCNQ2 from PV-INs could improve RTG’s anti-seizure efficacy in PTX-induced seizures (drug × genotype interaction on seizure latency: F=8.34, *p*=0.006).

**Figure 2.**
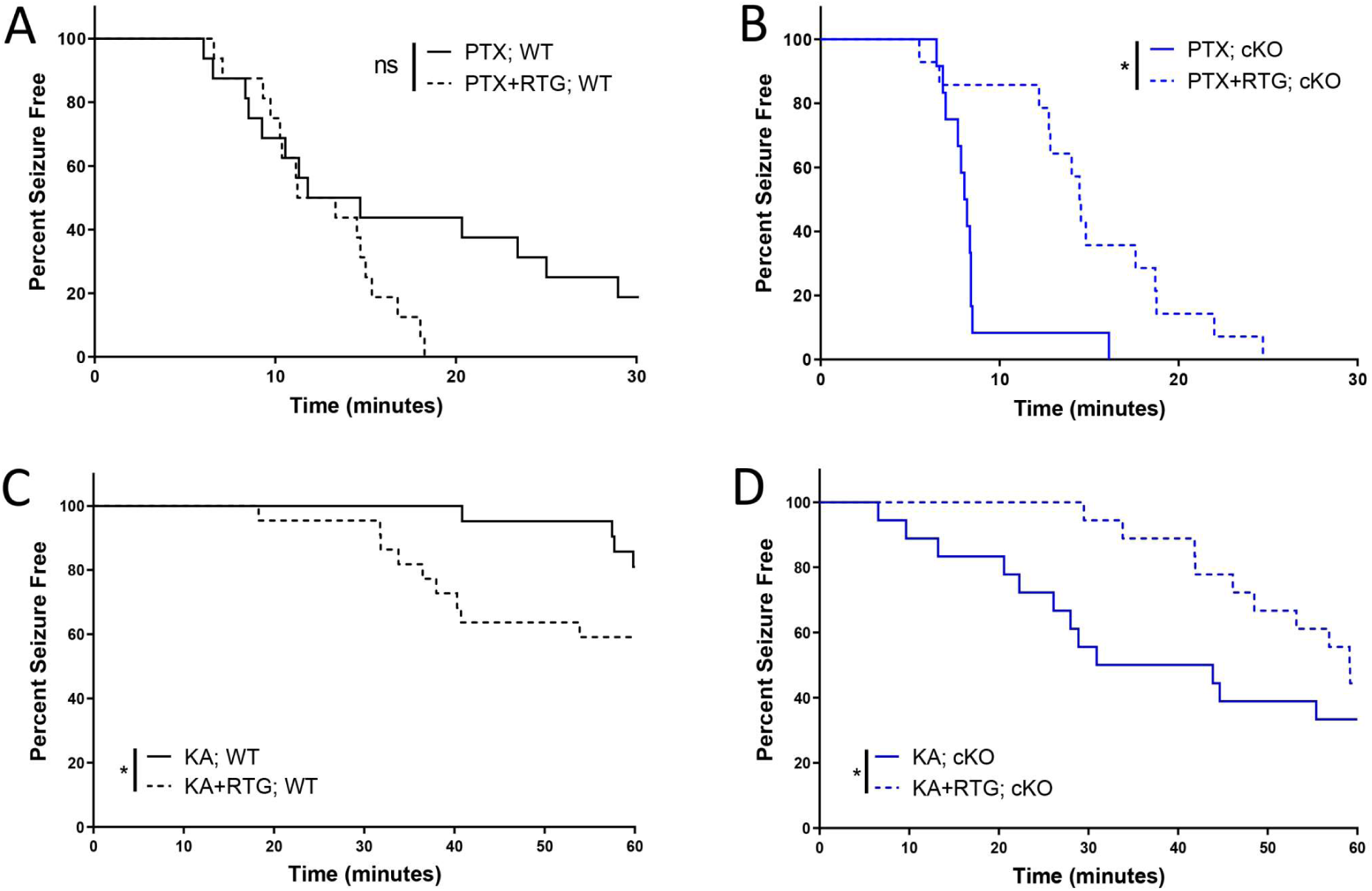
Improved efficacy of RTG in preventing PTX- and KA-induced seizures in cKO mice. Percentage of (A) WT (*p*=0.059; n=16, 16) and (B) cKO (*p*=0.001; n=12, 14) mice remaining seizure free with or without RTG plotted against time after PTX injection (Gehan-Breslow test). Percentage of (C) WT (*p*=0.042; *n*=21, 22) and (D) cKO (*p*=0.040; *n*=18, 18) mice with or without RTG administration plotted against time after KA injection (Grehan-Breslow test).

We then used intraperitoneal kainic acid (KA), another chemoconvulsant with a different induction mechanism, to test if the improved efficacy of RTG in cKO mice was independent of the seizure model. Aligned with findings from the PTX model, cKO mice had a significantly lower threshold to KA-induced seizures (p<0.001, Gehan-Breslow Test). In WT mice, administration of RTG (10mg/kg, i.p.) 15 minutes prior to KA injection (30mg/kg, i.p.) resulted in a significantly shorter latency to seizure onset compared to vehicle-injected mice (*p*<0.05, Grehan-Breslow test, Fig. 2C). In contrast, RTG injection in cKO mice significantly lengthened the seizure latency compared to vehicle-injected controls (p=0.040, Fig. 2D), indicating that conditional KO could improve RTG’s anti-seizure efficacy in KA-induced seizures as well (drug x genotype interaction on seizure onset: F=15.71, p<0.001). Of note, PTX- and KA-induced mortality was not significantly different with RTG treatment in either genotype, but there was a trend toward improvement in cKO mice (Fig S1).

### Selective loss of KCNQ2 and KCNQ3 expression at AISs of PV-INs in cKO mice

To confirm KCNQ2 was specifically lost from PV-INs in cKO mice, we performed immunostaining for KCNQ2 and KCNQ3 in the hippocampus and neocortex and imaged their signals in the neurons with wide-field and confocal microscopy. We also simultaneously labeled the axon initial segment (AIS) of PV-INs with AnkG immunostaining (see Methods). Labeling of AIS in PV-INs was achieved with the least ambiguity in stratum oriens and stratum radiatum in the hippocampal CA1 (Fig. 3A, B). In WT mice, CA1 PV-INs had KCNQ2 immunostaining particularly enriched in the distal AIS, overlapping with AnkG (Fig. 3A, C-E). In cKO mice, CA1 PV-INs showed no detectable KCNQ2 signals, consistent with the selective removal of KCNQ2 from PV-INs in these transgenic mice (Fig. 3B, F-H).

**Figure 3.**
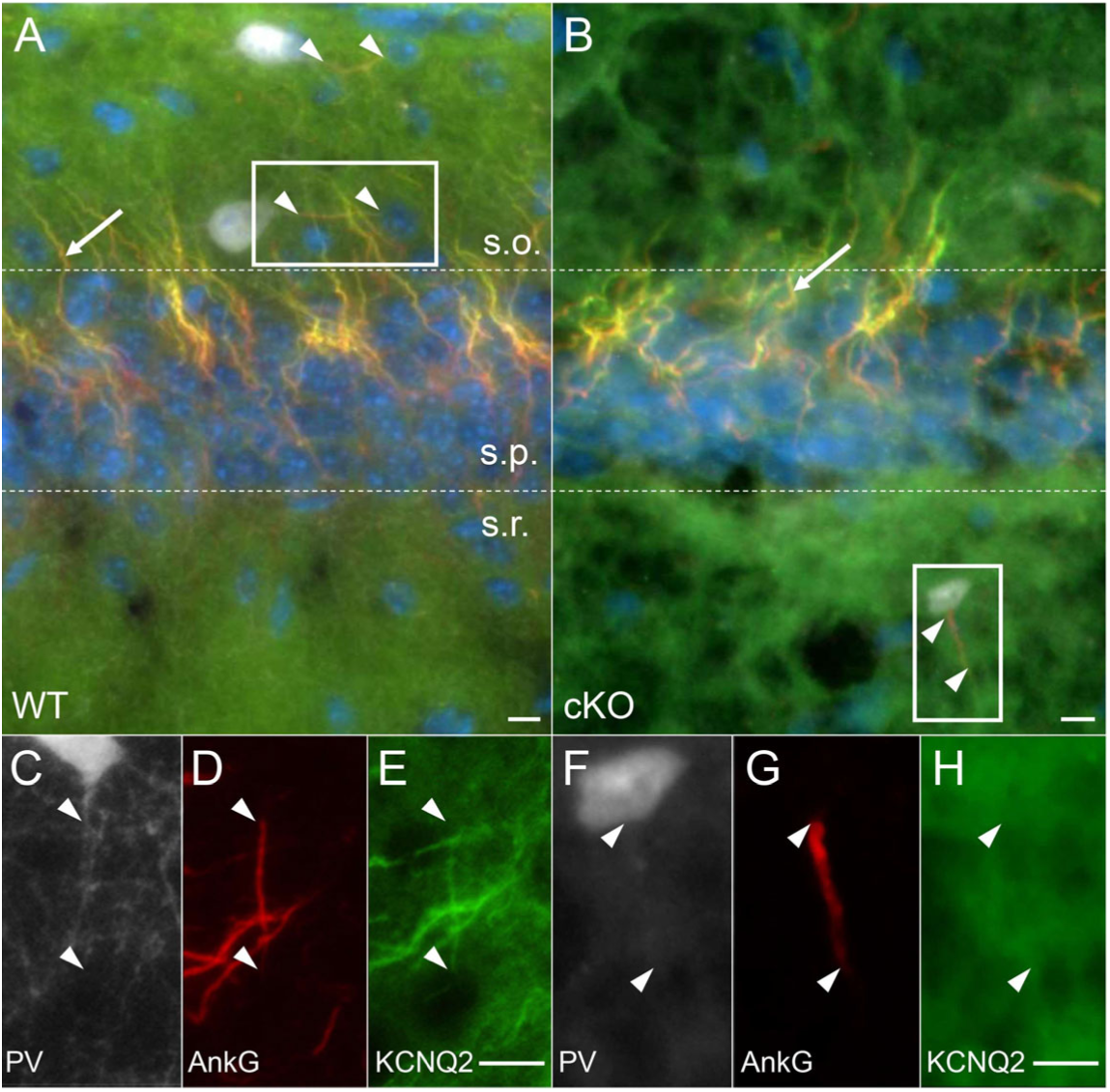
Selective deletion of KCNQ2 from CA 1 PV-IN in cKO mice. (A) In WT mice, KCNQ2 (green) was co-expressed with AnkG (red, arrowhead) at the distal axon initial segment (AIS) of PV-IN (TdTomato – white) in stratum oriens (s.o. = stratum oriens, s.p. = stratum pyramidale, s.r. = stratum radiatum). (B) In cKO mice, KCNQ2 was no longer co-expressed with AnkG in PV-IN (here shown in stratum radiatum of CA1), but remained evident in the AIS of CA1 pyramidal cells in stratum pyramidale (arrow). (C-E) Inset from A, showing KCNQ2 expression in the distal 2/3 of the AIS. (F-H) Inset from B, showing absence of KCNQ2 immunoreactivity localized to the PV interneuron AIS (arrowheads showing beginning and end of AIS). Scales: 10 µm.

KCNQ2 immunostaining was also particularly enriched at the distal AIS in WT PV-INs in the neocortex (Fig. S2A-D), and absent from PV-INs in cKO mice (Fig. S2E-H). In WT mice (n=4), 175 out of 177 (99.7±0.003%, mean±SEM) AISs (positive for AnkG) counted from non-PV-INs were positive for KCNQ2 (KCNQ2^+^). In PV-INs, all of the counted AISs (9 out of 9; 100%) were KCNQ2^+^. In cKO mice (n=5), almost all AISs (271 of 277; 98.3±0.008%) counted from non-PV-INs remained KCNQ2^+^, but all AISs from PV-INs were devoid of KCNQ2 (0%; 0/15), confirming the selective abolishment of KCNQ2 expression in PV-INs in cKO mice.

Given the extensive colocalization of KCNQ2 and KCNQ3 (Pan et al., 2006; Soh et al., 2014) and the sensitivity of KCNQ3-containing channels to RTG (Wang et al., 2018), we also imaged the AISs of PV-INs from WT and cKO mice after immunostaining for KCNQ3. In WT, KCNQ3 immunostaining was observed in the distal AISs of both CA1 PV-INs and non-PV-INs neurons (n=4, Fig. 4A-D). In cKO mice, KCNQ3 immunostaining was absent in PV-IN AISs (n=4, Fig. 4E-H), but remained detectable in neighboring non-PV-INs neurons, which were most likely CA1 pyramidal neurons (Fig. 4E-H). This suggests that the KCNQ2 and KCNQ3 subunits at WT PV-IN AISs form heteromeric channels, and that the KCNQ3 ankyrin-binding domain, though sufficient to target KCNQ3 to the AIS in vitro (Pan et al., 2006; Rasmussen et al., 2007), lacks this ability in the absence of KCNQ2 subunits in vivo.

**Figure 4.**
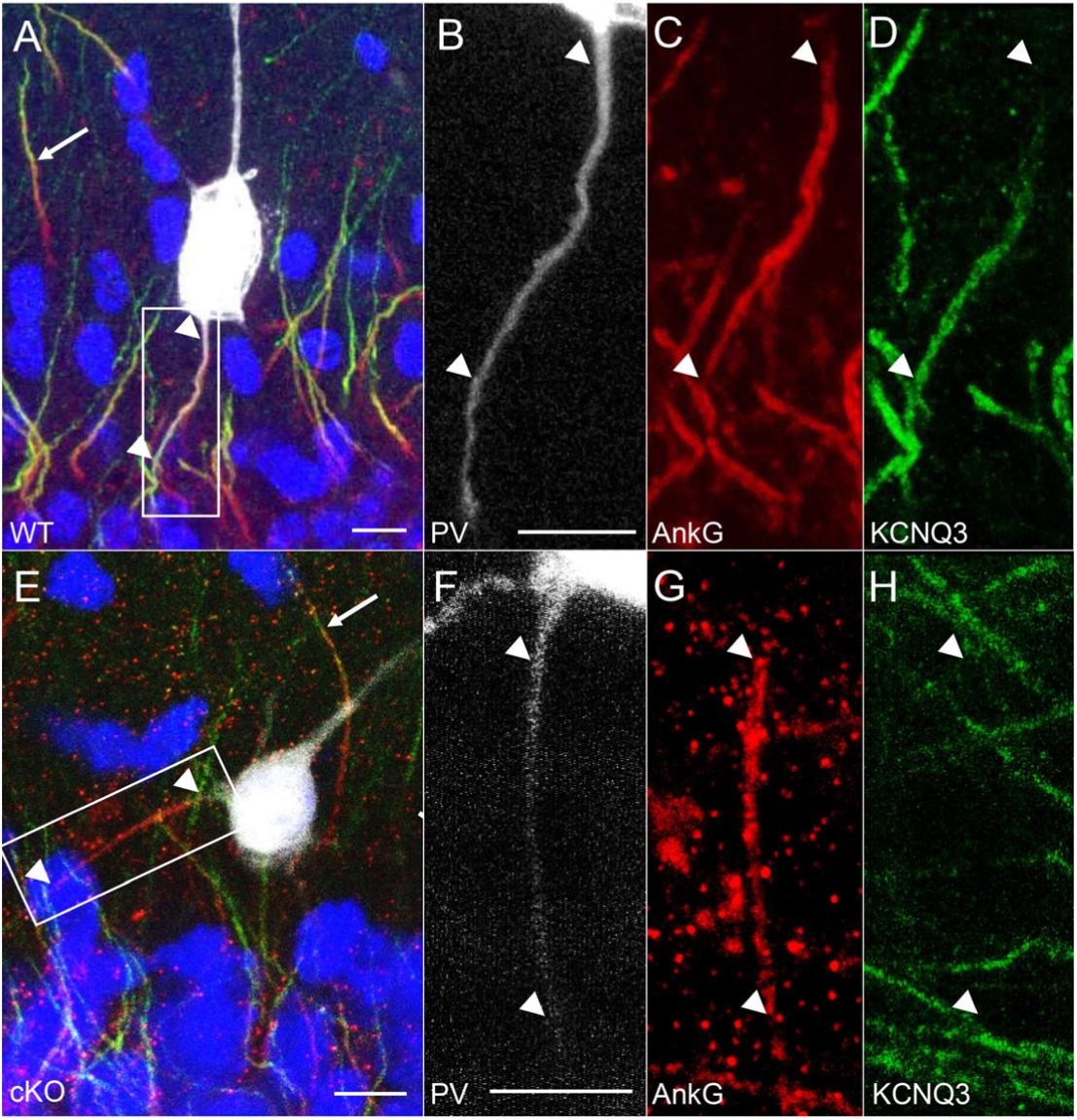
Absence of KCNQ3 AIS labeling in PV-INs in cKO mice. (A-D) With confocal imaging, KCNQ3 (green) expression was seen at the distal AIS (arrowheads) labeled by AnkG (red) in a WT CA1 stratum pyramidale PV-IN, with non-PV-INs also showing distal AIS labeling of KCNQ3 in the opposite direction (arrow). (E-H) In a PV-IN from the cKO mouse, staining for KCNQ3 was absent in the AIS (arrowheads), but detected in the neighboring AIS of a non-PV-IN (arrow). Scale bars: 10 µm.

### Decrease of M-currents and their sensitivity to RTG on PV-INs in cKO mice

KCNQ2/3-containing potassium channels are the major Kv7 channels mediating M-current (*I*M) in hippocampal neurons (Shah et al., 2002; Lawrence et al., 2006; Soh et al., 2014). Given our immunostaining results shown above, we expected that the M-currents in PV-INs would be significantly reduced in cKO mice, and the enhancing effect of RTG, an ASD known to promote the open state of KCNQ2/3-containing Kv7 channel (Main et al., 2000; Gunthorpe et al., 2012), on the M-currents would be blunted in cKO mice as well. To confirm these, we performed patch-clamp recordings on CA1 PV-INs to isolate the *I*_M_ from these neurons. In WT PV-INs, the amplitude of *I*_M_ at the baseline condition was 12.2±1.9 pA, which could be significantly enhanced by the application of RTG at 10 µM (30.9±3.9 pA, *p*<0.001, n=12 from three mice). RTG also increased the holding current with the voltage clamped at −20 mV in WT PV-INs (Fig. 5B; prior: 606±65pA; post: 716±69pA; *p*< 0.01, n=12). In cKO PV-INs, the amplitude of *I*_M_ induced with the same protocol was significantly smaller than WT (7.7±1.0 pA; *p=*0.03, n=12 from three mice), consistent with our previous study (Soh et al., 2014). Importantly, the *I*_M_ in cKO mice lost the sensitivity to 10 µM RTG (Fig. 5A, C, right, n=12 from three mice). There was a significant drug × genotype interaction on *I*_M_ current (F=24.1, *p*< 0.01). In addition, RTG had a smaller effect on *I*_hold_ from cKO PV-INs compared to WT, and there was a significant drug × genotype interaction on *I*_hold_ (Fig. 5B; F =6.9, p< 0.05). All these results confirmed that the current component carried by KCNQ2/3-containing channels was significantly decreased by gene knockout.

**Figure 5.**
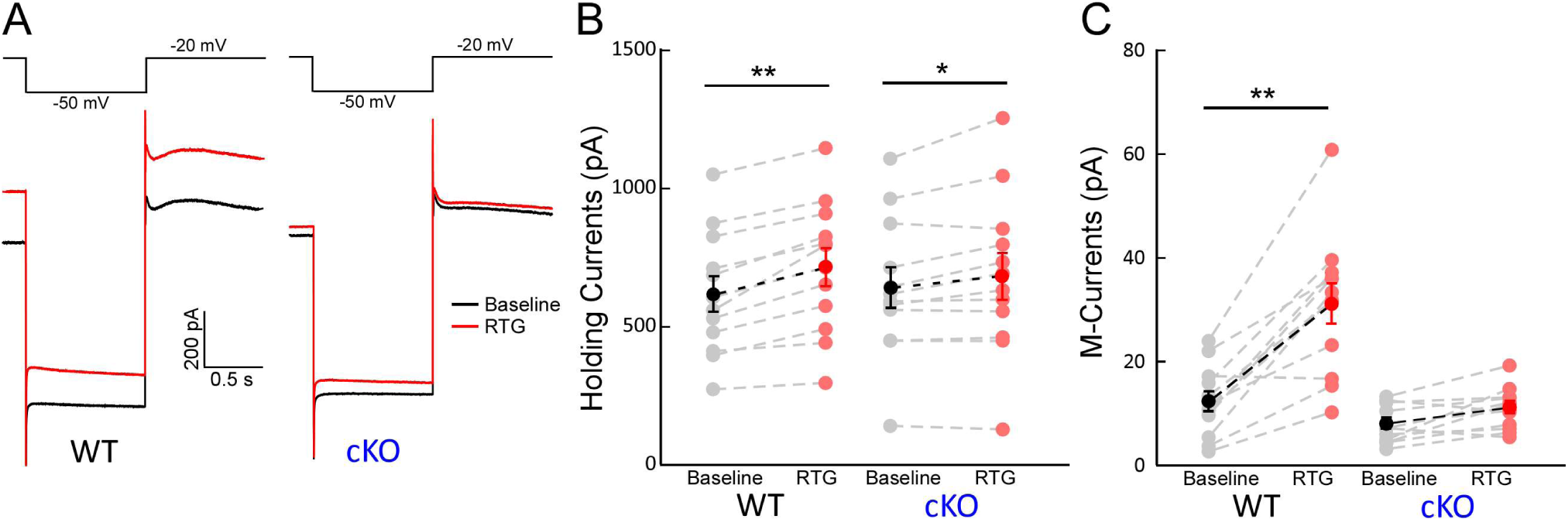
Effects of retigabine (RTG) on M-currents of hippocampal PV-INs. (**A**) Representative traces of M-currents (*I*M) in PV-INs from WT (left) and from cKO (right) before (black) and after (red) 10 µM RTG application. (**B**) RTG significantly increased the holding currents (*I*hold) at −20 mV in PV-INs from WT (left, n = 12) mice and cKO mice (right, n = 12), but RTG had less effects on *I*hold from cKO mice (\**p* < 0.05, \*\**p*< 0.01; interaction: F(1,1) =6.9, p< 0.05 with two-way ANOVA). (**C**) RTG significantly increased the *I*M amplitude in PV-INs from WT mice, but not in PV-INs from cKO mice (WT, n = 12; cKO, n = 12; ** *p*< 0.01; interaction: F(1,1)=24.1, p< 0.01 with two-way ANOVA).

### Suppression of PV-INs by RTG was selectively blunted in cKO mice

Given the above results, we reasoned that RTG could suppress the excitability of WT PV-INs, but this suppressive effect was selectively blunted in these neurons from cKO mice (Grigorov et al., 2014; Paz et al., 2018). We thus performed patch-clamp recordings on both CA1 pyramidal cells (PC) and PV-INs (Fig. 6) to test if there was any selective change for PV-INs in their responses to RTG in cKO mice compared to PCs. Cell responses to depolarizing current steps were recorded, and the number of action potentials (APs) was plotted as a function of the injected current (AP-I curve; Fig. 6B-C). Consistent with our previous findings (Soh et al., 2018), the AP-I curve of PV-INs in cKO shifted left compared to WT when injected currents were increasing, indicating the enhanced excitability of PV-INs in the absence of *Kcnq2* (Fig. 6D; n=19 PV-INs from 4 WT mice; n=18 PV-INs from 4 cKO mice). AP threshold for PV-INs was not significantly different between the two genotypes (Suppl. Table, *p*>0.05), suggesting Kv7 channels exert no prominent effect on AP threshold. Bath application of RTG (10 µM) for 10 min significantly reduced the number of APs in response to the suprathreshold current steps in WT PV-INs (Fig. 6C, *p*<0.01). In cKO PV-INs, while RTG still suppressed the number of APs in response to some current injections, the suppressive effect of RTG was significantly smaller in cKO mice (Fig. 6C, gene x drug interaction: F=4.3, *p*< 0.05). Importantly, as expected in CA1-PCs, there were no significant differences in AP-I curves between WT and cKO (Fig. S3, n=19 PCs from 3 WT mice; n=21 PCs from 3 cKO mice), and no significant difference in RTG-induced suppression of AP frequency (Fig. S3A-C).

**Figure 6.**
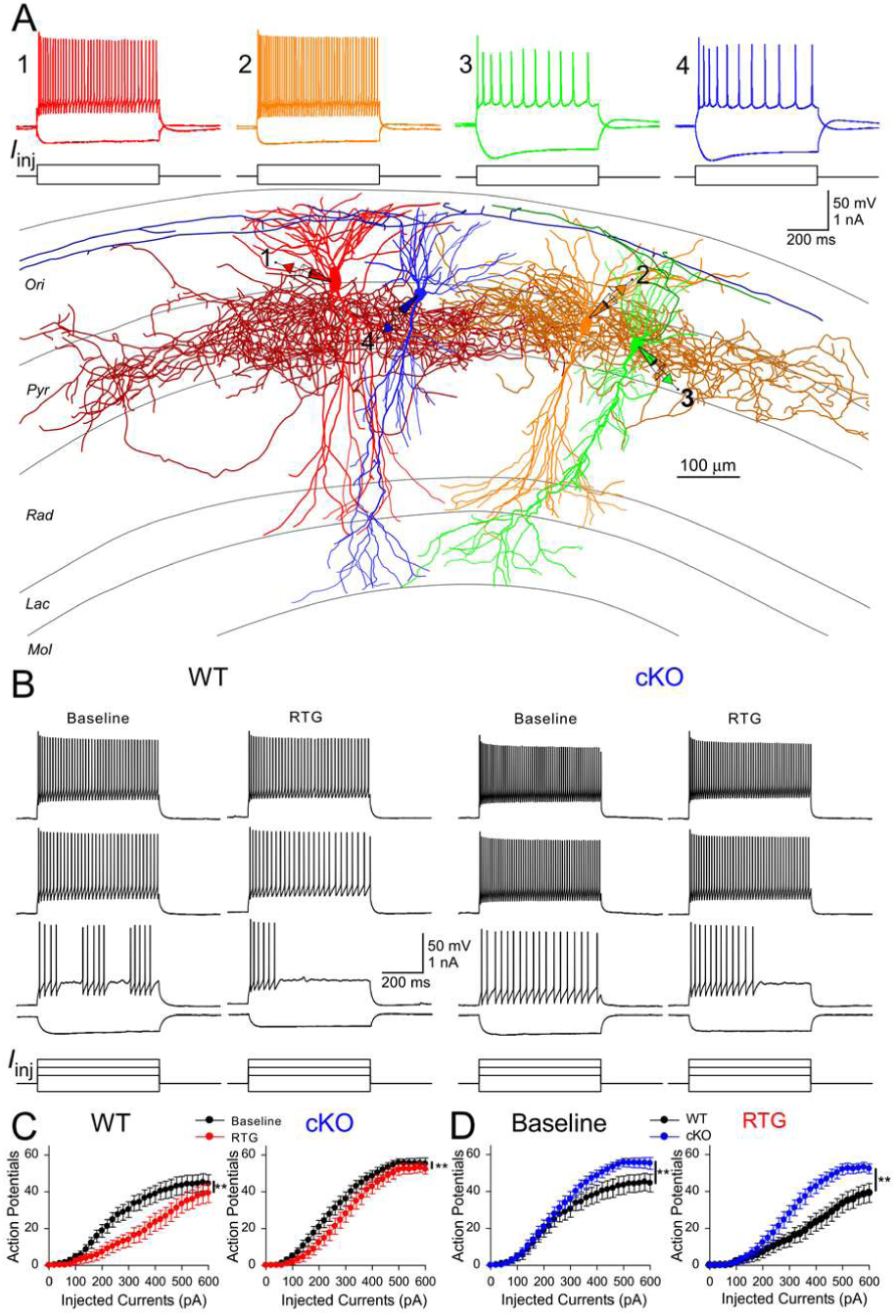
Effects of retigabine (RTG) on the excitability of PV-INs in vitro. (**A**) Experimental configuration in CA1. Whole-cell recordings were obtained simultaneously from four neurons, including two PV-INs (cell 1 and 2) and two PCs (cell 3 and 4). PV-INs were distinguished from PCs and other interneurons by firing pattern (above) and post hoc morphology recovery (below). (**B**) Representative membrane voltage responses to different current step injections in PV-INs from WT (left) and cKO (right) before and after RTG application. (**C**) RTG suppressed APs in PV-INs from WT (left, n = 19, \*\**p*< 0.01 with two-way ANOVA) and cKO mice (right, n=18, \*\**p*< 0.01 with two-way ANOVA), but RTG had less suppressive effects on PV-INs excitability from cKO mice (gene x drug interaction: F=4.3 \*\**p*< 0.05 with two-way ANOVA). (**D**) Left: at the baseline, PV-INs fired more APs in cKO mice when injection currents were more than 250 pA (F=4.4, *p*< 0.05 with two-way ANOVA). Right: In the presence of RTG, PV-INs fires more APs in cKO mice as well across a large range of current injections than WT mice (F=8.5, \*\**p*< 0.01 with two-way ANOVA).

We also examined the change of RTG-induced suppression of PV-INs in cKO mice using a ramp stimulation protocol (Varghese et al., 2021). Kv7 channels have slow activation kinetics; thus, slow ramp protocols reveal different aspects of the roles of Kv7 channels in regulating neuronal excitability. As observed with our step protocols, retigabine application reduced the number of APs during the slow ramp protocol in WT PV-INs (Baseline: 88.2 ± 9.7; RTG: 26.0 ± 6.6, n = 19; *p*< 0.01, paired Student’s t test). While RTG application still suppressed the number of APs in cKO PV-INs (Fig. 7A, B; Baseline: 102.9±10.0; RTG: 72.9± 10.2, n = 19; *p*< 0.01, paired Student’s t test), the suppressive effect was significantly smaller in cKO mice (Genotype x Treatment interaction: F=9.5, *p*<0.01). These results indicate the reduced suppression of PV-INs by RTG in cKO mice is independent of stimulation protocol.

**Figure 7.**
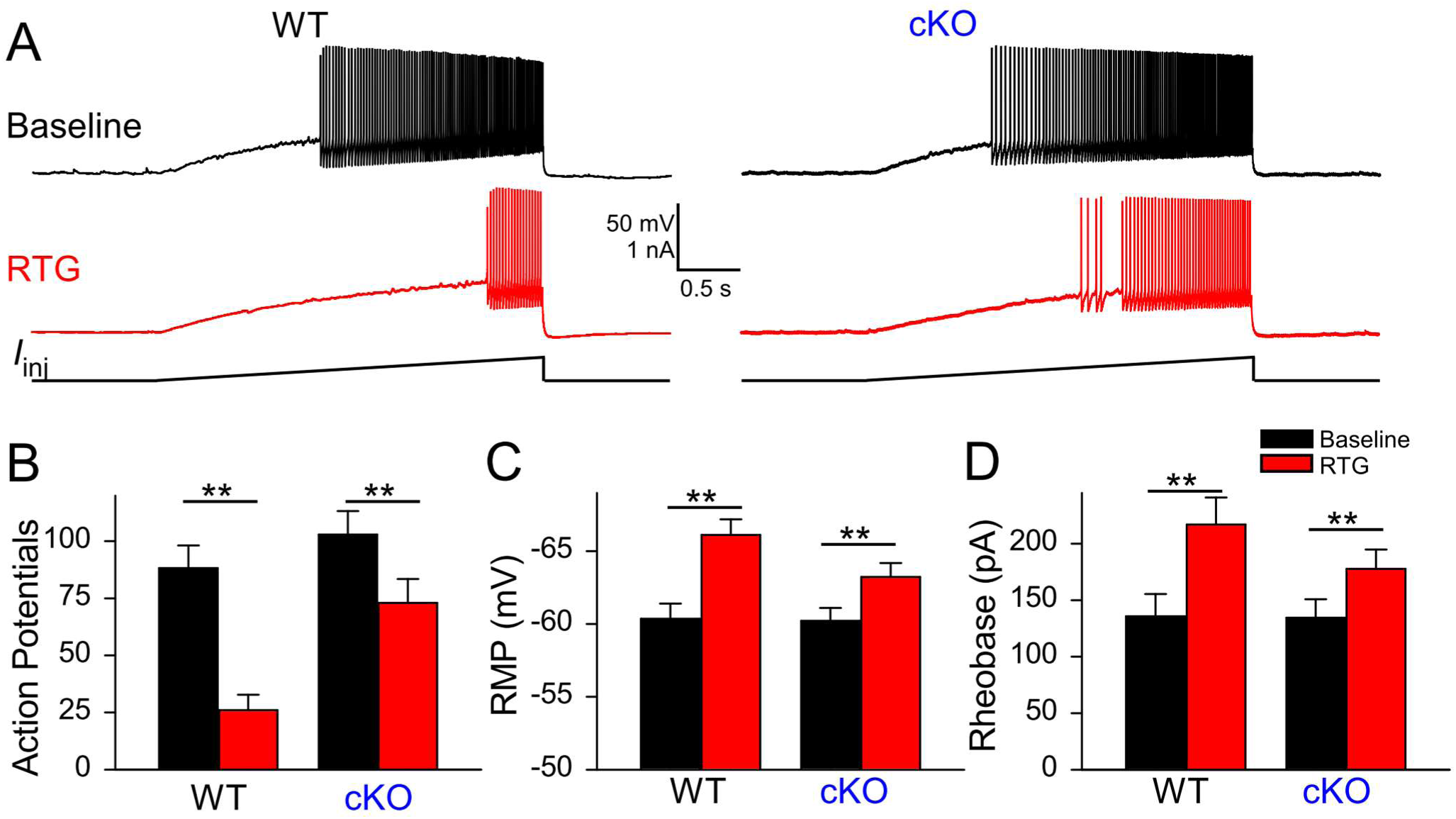
Retigabine’s effect on excitability of cKO PV-INs. **A:** representative voltage responses to a ramp protocol in CA1 PV-INs from WT and cKO slices before and after bath application of RTG (10 µM). (**B)**: summary graphs showing the effect of 10 µM retigabine (RTG) on action potential number (WT, n = 19; cKO, n = 19; \*\**p*< 0.01, Genotype x Treatment interaction: F=9.5, *p*<0.01 with two-way ANOVA). (**C**) Bath application of RTG in hippocampal slices significantly hyperpolarized the RMP of PV-INs from both WT and cKO mice, but RTG had a smaller effect on RMP from cKO mice (WT, n = 19; cKO, n = 18; \*\**p*< 0.01; interaction: *p*= 0.046 with two-way ANOVA). (D) Bath application of RTG significantly increased the rheobase of PV-INs from both WT and cKO mice, but RTG had less effects on cKO mice (WT, n = 19; cKO, n = 18; \*\**p*< 0.01, interaction: p= 0.046 with two-way ANOVA).

Given that Kv7 channel can regulate intrinsic membrane properties, we also examined the effect of RTG on the passive membrane properties of CA1-PCs and PV-INs from both genotypes. As expected, 10 µM RTG significantly hyperpolarized the resting membrane potential (RMP) of PV-INs from WT mice (Fig. 7C and Suppl. Table; *p*<0.01, Wilcoxon test; n=19 PV-INs from 4 WT mice). RTG could induce the hyperpolarization of PV-INs in cKO mice as well (7C and Suppl. Table; *p*<0.01, Wilcoxon test; n= 18 from 4 cKO mice), but the effect was significantly smaller in cKO mice (drug × genotype interaction on RMP: F=4.3, *p*<0.05). There was no significant difference in the rheobase of PV-INs between WT and cKO (Fig. 7D, Suppl. Table). RTG significantly increased the rheobase of WT PV-INs (Fig. 7D, *p*<0.01), but this effect became smaller in cKO PV-INs (Fig. 7D and Suppl Table, drug × genotype interaction: F=4.3, p<0.05). The effect of RTG on other parameters in WT and cKO are detailed in Suppl. Table. Of note, RTG could increase the AP amplitude and reduce AP width in WT PV-INs, but these effects were significantly decreased in cKO mice, indicating that KCNQ2/3-containing Kv7 channels could modulate the AP shape of PV-INs. Together, these findings indicated that RTG suppressed the excitability of PV-INs by modulating their intrinsic membrane properties, and these effects were significantly reduced in cKO mice.

In CA1-PCs, RTG application significantly decreased their input resistance, and this effect was not significantly changed in cKO mice (Suppl. Table; n=19 PCs from 3 WT mice; n=21 PCs from 3 cKO mice). Surprisingly, different from its effect on PV-INs, RTG could not hyperpolarize the RMP of CA1-PCs from either WT mice or cKO mice, suggesting the Kv7 channels activated by RTG in CA1-PCs has little effect on the RMP of these neurons. RTG application increased the rheobase of CA1-PCs, and this effect was significantly enhanced in cKO mice (Suppl. Table), likely due to a secondary effect of selective gene removal (Soh et al., 2018). The effect of RTG on other parameters in WT and cKO was shown in Suppl. Table, and there were no significant differences in these parameters between WT and cKO.

## Discussion

ASD targets are generally expressed in both excitatory neurons and GABAergic inhibitory interneurons (INs). Therefore, while ASDs suppress the excitability of principal excitatory neurons, they may simultaneously suppress IN excitability. The simultaneous actions on INs and excitatory neurons may reduce their anti-seizure efficacy. However, this hypothesis has not previously been directly tested. The role of INs in seizures has been controversial, with some studies indicating that IN activity can suppress seizures (Krook-Magnuson et al., 2013; Lu et al., 2016; Călin et al., 2018), support seizures (Khoshkhoo et al., 2017; Miri et al., 2018), or both (Assaf and Schiller, 2016; Magloire et al., 2019), depending on the paradigm. To further illuminate this issue, we generated cKO mice to remove KCNQ2 from PV-INs and compared the efficacy of retigabine (RTG), an agonist for KCNQ2-containing Kv7 channels, in WT and cKO mice *in vivo* and *in vitro*. We found that removal of KCNQ2 from PV-INs resulted in the additional removal of KCNQ3, selectively blunted RTG-induced suppression of PV-INs, and improved the anti-seizure efficacy of RTG in two chemoconvulsive seizure models.

### Improved anti-seizure efficacy of RTG in cKO mice

WT and cKO mice had equivalent growth curves, behavior, survival, and baseline EEG spectra, removing some potential study confounds. In WT mice, 10 mg/kg RTG was ineffective in delaying the onset of seizures when challenged with 10 mg/kg PTX (Fig. 2). In a prior study using 2.5 mg/kg PTX, RTG could prevent PTX-induced seizures in a different strain of WT mice, with an ED_50_ of 18.6 mg/kg (Rostock et al., 1996). Therefore, the lack of RTG efficacy in our WT mice may be due to a relatively low dose of RTG, a relatively high dose of PTX, or strain differences (Frankel, 2009). cKO mice had a significantly lower seizure threshold with PTX challenge, as seen previously (Soh et al., 2018), but RTG significantly delayed seizure onset in these mice, indicating the anti-seizure efficacy of RTG could be improved with conditional removal of *Kcnq2* from PV-INs.

Given that PV-INs exert influence on network activity primarily via GABA_A_ receptors, the receptors that PTX blocks, a potential confound could arise from using the PTX seizure model to evaluate the effect of a drug on PV-INs. Therefore, we also tested our hypothesis using an acute kainic acid (KA) seizure model. Interestingly, RTG significantly hastened seizure onset in WT mice, similar to previous findings in rats (Friedman et al., 2015). In cKO mice, this paradoxical seizure aggravation with RTG was not only rescued, but reversed. These results suggest that RTG-indued seizure aggravation in WT mice may be due to its simultaneous suppression of inhibitory PV-INs, and selectively blunting this suppression (as seen in vitro data) results in improved anti-seizure efficacy. Seizure aggravation after ASD treatment has been well-described in mouse models of Dravet Syndrome, which carry mutations in the *Scn1a* sodium channel subunit with notable PV-IN dysfunction (Yu et al., 2006; Ogiwara et al., 2007). ASDs that block sodium channels exacerbate seizures in both patients and mice with this mutation, presumably due to their simultaneous suppression of PV-IN activity. In addition, in the *stargazer* mouse model of absence epilepsy, we find ASD-induced paradoxical seizure exacerbation to be linked to its simultaneous suppression of inhibitory neurons (Maheshwari et al., 2013). These results suggest that simultaneous suppression of INs by an ASD reduces its anti-seizure efficacy, while sparing inhibitory neurons could improve anti-seizure efficacy.

### Subcellular location of KCNQ2/3 in the cortex

In this study, we found that the expression of both KCNQ2 and KCNQ3 were specifically enriched in the AIS of PV-INs in the hippocampal CA1 region and neocortex. Deletion of *Kcnq2* from PV-INs removed KCNQ2 from the AIS, but it also removed KCNQ3 from PV-INs, consistent with our previous finding in pyramidal cells (PC) (Soh et al., 2014). Further investigation may be needed to understand the mechanism underlying the subsequent loss of KCNQ3 subunits upon removal of KCNQ2 and whether compensatory mechanisms are present at different stages of development (Carver and Shapiro, 2019). Our findings of specific AIS location of Kv7 channels in PV-INs corroborate and extend previous findings in neocortical PCs (Battefeld et al., 2014; Soh et al., 2014), highlighting the highly compartmentalized functions of Kv7 across cell types. Nevertheless, it may not be the case that all cell types have such a highly restricted Kv7 expression. For instance, hippocampal CA1 somatostatin-expressing interneurons appear to have a high expression of KCNQ2/3 in the soma with prominent somatic M-currents, in addition to having these channels highly expressed in the AIS (Cooper et al., 2001; Lawrence et al., 2006).

### Decreased effect of RTG on PV-INs in cKO mice

With the selective removal of KCNQ2/3 from PV-INs, M-current (*I*_M_) in PV-INs, the current mediated by Kv7 channels, was significantly reduced in cKO mice. In addition, the enhancing effect of RTG, an ASD known to promote KCNQ2/3-containing Kv7 channel, on the M-currents was blunted, consistent with our immunostaining results. As expected, RTG reduced APs of PV-INs elicited by depolarizing current steps, reduced their input resistance, hyperpolarized their RMP, and increased their rheobase. Similar effects have been observed in somatosensory L5 PCs upon the activation of KCNQ2/3-containing potassium channels (Battefeld et al., 2014). Interestingly, while RTG could also dampen the spiking activity of CA1-PCs, RTG did not induce any significant change in somatic RMPs of CA1-PCs, in contrast to what we observed in PV-INs, but consistent with previous studies on CA1-PCs (Soh et al., 2014). This cell type-specific difference may be due to a more hyperpolarized RMP of CA1-PCs than PV-INs and neocortical L5 PCs. Under our recording conditions, CA1-PCs have an RMP that is near the foot of the voltage-gated activation curve for Kv7 channels, and the fraction of somatic Kv7 channels that are active at the RMP would thus be lower in PCs compared with PV-INs.

RTG also lost most of the capability to suppress APs elicited by different protocols (Fig. 6C, Fig. 7), while its suppressive effect on CA-PCs remained unaffected (Fig. S3C). RTG still retained a small suppressive effect in cKO PV-INs, and this RTG-sensitive, residual component may be related to the presence of KCNQ5 homotetramers (Fidzinski et al., 2015), or non-Kv7, “off-target” effects of RTG (Otto et al., 2002; Treven et al., 2015). In addition, RTG increased the rheobase and hyperpolarized RMPs of WT PV-INs, but these effects were significantly decreased in cKO PV-INs. Nevertheless, the cKO did not significantly impact the effect of RTG on the input resistance of PV-INs (Suppl. Table, p>0.05), suggesting the RTG-induced change in input resistance may not be mediated by KCNQ2/3-containing Kv7 channels. The effects of cKO on PV-INs were not observed in CA1-PCs from cKO mice, except with the rheobase of CA1-PCs (Suppl. Table), which may be due to a secondary effect of selective gene removal (Soh et al., 2018).

### Sparing interneurons to improve anti-seizure efficacy

In summary, the anti-seizure efficacy of RTG was improved in a model-independent manner when its drug target was removed from a major group of INs. RTG lost most of the capability to suppress the excitability of these INs in a protocol-independent manner, while its suppressive effects on excitatory neurons remained intact. We reasoned the improved anti-seizure efficacy of RTG seen in vivo was due to reduced suppression of INs by RTG. Therefore, sparing interneurons by ASDs may be beneficial for anti-seizure efficacy. Developing ASDs that have less effect on INs thus may be an effective strategy to improve the efficacy of ASDs. For instance, innovative approaches such as DART (drugs acutely restricted by tethering) could be harnessed to alter the cell type-specificity of drug action to improve ASD efficacy (Shields et al., 2017).

There are some limitations to our study. First, *Kcnq2* deletion was cell-type restrictive but not developmentally restricted, and thus could change the network in ways that may affect seizure characteristics. Nevertheless, PTX- and KA-induced seizure semiologies of cKO and WT mice were similar, including behavioral progression to characteristic Racine Stage 5 generalized convulsions (Soh et al., 2018). In addition, our studies were performed on chemoconvulsant-induced seizures, and validation of our findings in models of spontaneous, recurrent, and pharmacoresistant seizures would enhance the generalizability of these findings. Finally, the current focus was on PV-INs, but further study should include exploring how the changes in drug effects on PCs and other interneuron subtypes impact their anti-seizure efficacy (Tremblay et al., 2016).

Our findings may also have important implications for ASD pharmacoresistance, affecting one in three patients with epilepsy (Kwan and Brodie, 2000; Tang et al., 2017). The target hypothesis of pharmacoresistance suggests that drug effect is limited by altered expression of its targets (Pohlmann-Eden and Weaver, 2013), while the intrinsic severity hypothesis posits that pharmacoresistance develops as the underlying disease process becomes more severe (Rogawski, 2013). Despite the greater severity of seizures when challenged with chemoconvulsant, cKO mice responded more favorably to RTG than WT. Our findings thus support the target hypothesis over the intrinsic severity hypothesis. It will be interesting to see if this conclusion is sustained in a broader range of seizure models.

## Acknowledgments

We acknowledge funding from NINDS R01 NS101596 (AT), NINDS R01 NS49119 and the Jack Pribaz Foundation (ECC), NIMH R01 MH109556 (JJ, XJ), NIMH R01 MH120404 (JJ, XJ), and NINDS K08 NS096029 (AM). Research reported in this publication was also supported by P50HD103555 for use of the Microscopy Core facilities.

## Ethical Publication Statement

We confirm that we have read the Journal’s position on issues involved in ethical publication and affirm that this report is consistent with those guidelines.

---

**Figure S1.**
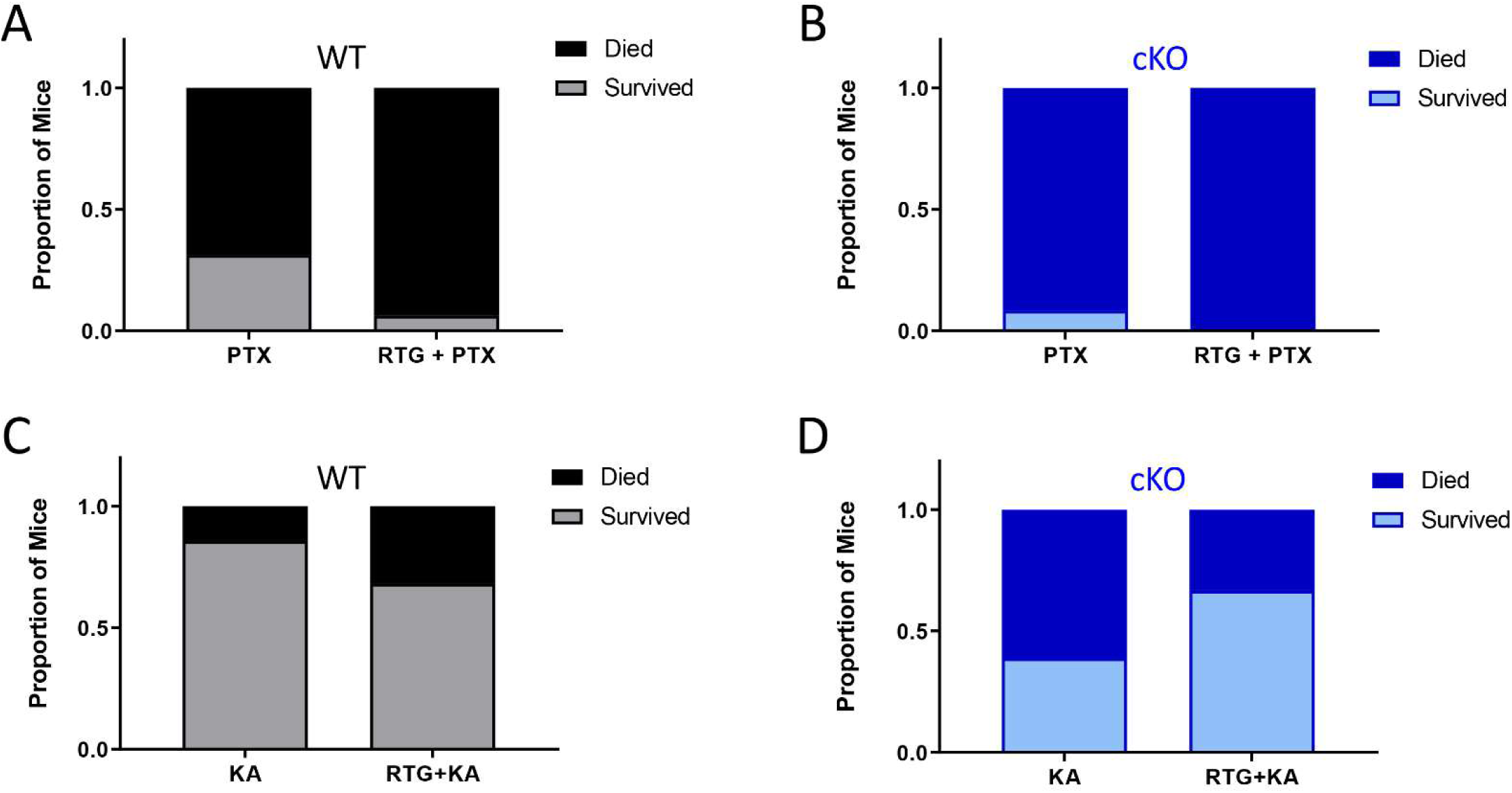
No change in seizure-induced mortality with RTG for cKO mice. Mortality following PTX-induced seizures was not significantly altered with RTG treatment in either (A) WT (n=16, 16) or (B) cKO (n=12, 14) mice. Mortality following KA-induced seizures was not significantly altered with RTG treatment in either (C) WT (n=21, 22) or (D) cKO (n=18, 18) mice.

**Figure S2.**
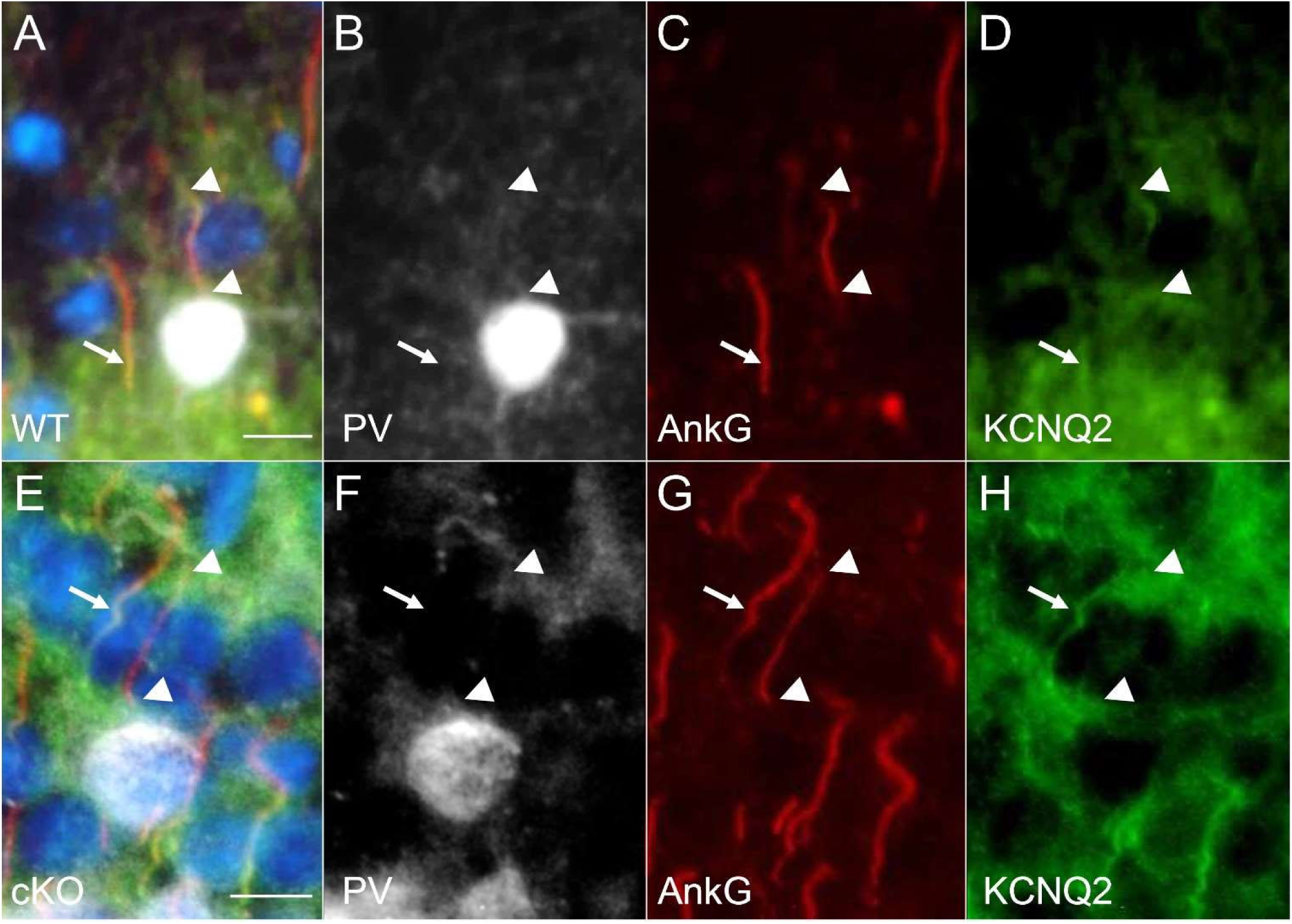
Selective deletion of KCNQ2 from cortical PV-IN in cKO mice. (A-D) In WT mice, KCNQ2 (green) was co-expressed with anti-AnkG (red) at the distal axon initial segment (AIS) of PV-IN (TdTomato – pseudocolored in white), seen here in Layer 2 of the neocortex (arrowheads) as well as non-PV-IN (arrow). (E-H) In cKO mice, KCNQ2 was no longer co-expressed with AnkG in PV-INs (arrowheads) but remained evident in the AIS of non-PV-INs (arrow); bar = 10 µm.

**Figure S3.**
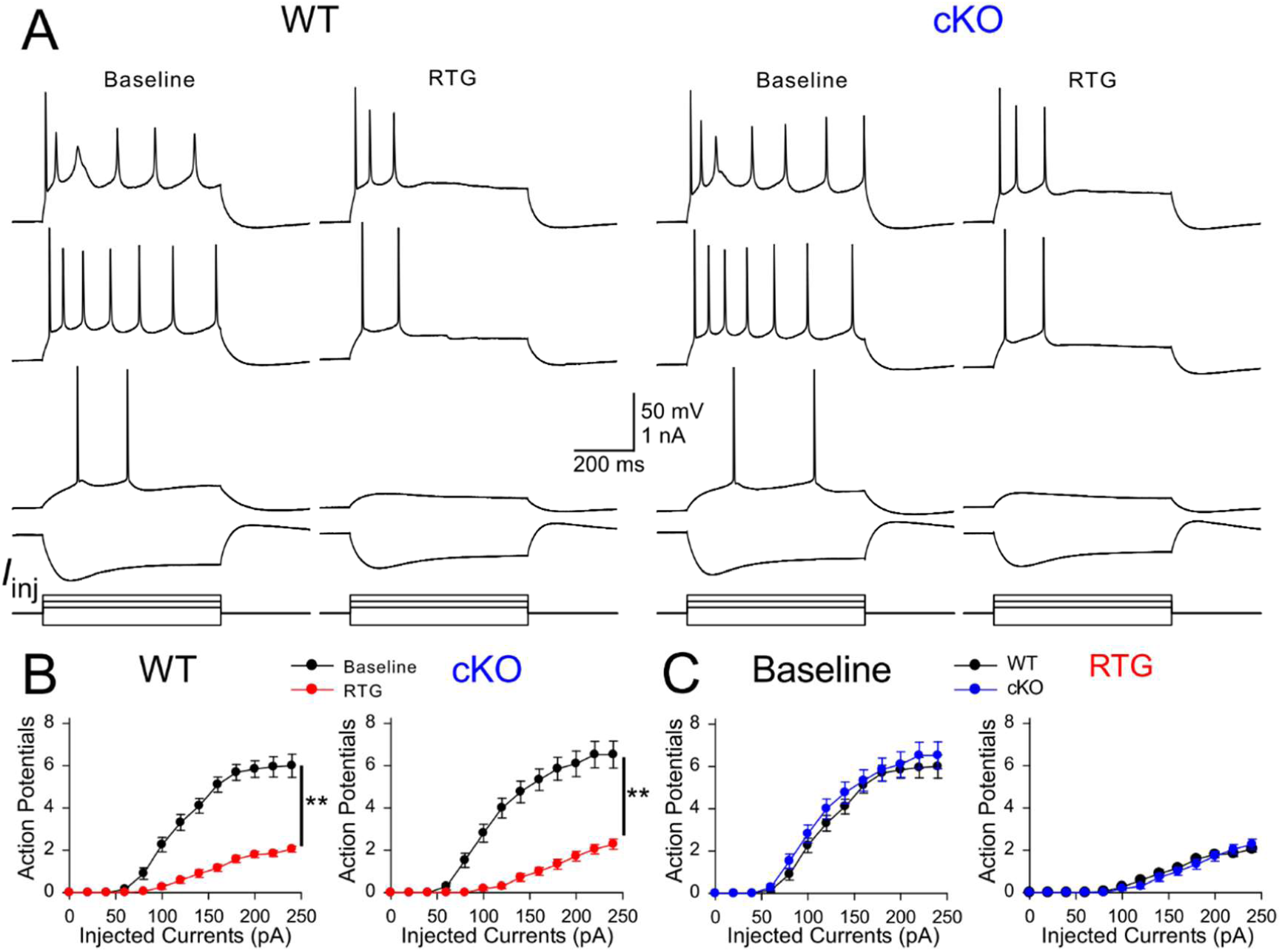
Effects of RTG on the excitability of hippocampal CA1-PCs *in vitro*. (A) Representative membrane voltage responses to different current steps in CA1-PCs from WT (left) and cKO (right) before and after 10 µM RTG treatment. (B) RTG was effective in suppressing APs in WT PCs (left, n=19, \*\**p*<0.01 with two-way ANOVA) and in cKO PCs (right, n=21, \*\**p*<0.01 with two-way ANOVA). (C) *Kcnq2* conditional knock-out from PV-INs did not significantly change AP-I curve of CA1-PCs at baseline (left). RTG induced a similar degree of suppression on the excitability of CA1-PCs from WT and cKO mice (right).

**Table:** Active and Passive Membrane Properties of CA1 PV-INs and CA1-PCs in WT and cKO.

| Cell Type | WT PV-INs |  | cKO PV-INs |  | CA1-PC in WT |  | cKO CA1-PC in cKO |  |
| --- | --- | --- | --- | --- | --- | --- | --- | --- |
| Treatment | Baseline | RTG | Baseline | RTG | Baseline | RTG | Baseline | RTG |
| Rheobase (pA) | 135.79 ± 19.68 | 216.84 ± 24.00 ** | 134.44 ± 16.31 | 177.78 ± 17.10 ** | 90.53 ± 4.15 | 133.68 ± 7.18 ** | 88.57 ± 5.08 | 152.38 ± 8.34 ** |
| Input Resistance (MΩ) | 159.36 ± 19.90 | 119.25 ± 14.35 ** | 162.54 ± 14.97 | 121.35 ± 10.48 ** | 144.85 ± 5.73 | 114.95 ± 5.02 ** | 143.80 ± 6.71 | 108.74 ± 5.56 ** |
| RMP (mV) | -60.36 ± 1.04 | -66.11 ± 1.07 ** | -60.21 ± 0.89 | -63.22 ± 0.96 ** | -70.34 ± 0.74 | -69.15 ± 0.56 | -69.44 ± 0.70 | -69.31 ± 0.76 |
| AP Threshold (mV) | -36.69 ± 0.86 | -37.80 ± 0.96 | -35.93 ± 0.80 | -35.86 ± 0.99 | -37.01 ± 1.73 | -36.34 ± 1.93 | -39.56 ± 1.83 | -38.89 ± 1.81 |
| AP Amplitude (mV) | 66.62 ± 1.45 | 69.70 ± 1.50 ** | 66.16 ± 1.14 | 65.66 ± 1.15 | 88.12 ± 2.43 | 85.43 ± 2.81 ** | 90.75 ± 1.80 | 88.65 ± 1.71 ** |
| AHP (mV) | 24.06 ± 0.83 | 21.83 ± 0.87 ** | 24.24 ± 0.60 | 22.08 ± 0.67 ** | 10.05 ± 1.66 | 10.09 ± 1.60 | 10.01 ± 1.21 | 9.94 ± 1.23 |
| AP Half width (ms) | 0.61 ± 0.02 | 0.57 ± 0.02 ** | 0.58 ± 0.02 | 0.56 ± 0.02 | 2.07 ± 0.04 | 1.99 ± 0.04 ** | 2.00 ± 0.05 | 1.93 ± 0.04 ** |

AHP: afterhyperpolarization, AP: action potential, RMP: resting membrane potential. \*\* $p < 0.01$ RTG vs Baseline with Two-way Repeated Measures ANOVA. Genotype X Treatment interaction on rheobase of PV-INs: $F = 4.30$ , $p < 0.05$ ; interaction on RMP of PV-INs: $F = 4.3$ , $p < 0.05$ ; interaction on AP amplitude of PV-INs: $F = 5.2$ , $p < 0.05$ ; interaction on AP half width of PV-INs: $F = 4.7$ , $p < 0.05$ ; interaction on rheobase of CA1-PCs, $F = 4.5$ , $p < 0.05$ .

## Notes

**Conflicts of Interest** AM received funding from NeuCyte, Inc to perform studies that were independent from this research. ECC serves as a consultant to Knopp Biosciences and Xenon Pharmaceuticals, which develops products related to the research being reported. The terms of this arrangement have been reviewed and approved by the Baylor College of Medicine in accordance with its policy on Financial Conflicts of Interests in Research. The remaining authors have no conflicts of interest.

### Competing Interest Statement

AM received funding from NeuCyte, Inc to perform studies that were independent from this research. ECC serves as a consultant to Knopp Biosciences and Xenon Pharmaceuticals, which develops products related to the research being reported. The terms of this arrangement have been reviewed and approved by the Baylor College of Medicine in accordance with its policy on Financial Conflicts of Interests in Research. The remaining authors have no conflicts of interest.

